# On phage adsorption to bacterial chains

**DOI:** 10.1101/2020.06.10.144675

**Authors:** R. S. Eriksen, N. Mitarai, K. Sneppen

## Abstract

Bacteria often arrange themselves in various spatial configurations which changes how they interact with their surroundings. In this paper, we investigate *in silico* how the structure of the bacterial arrangements influences the adsorption of bacteriophages. We quantify how the adsorption rate scales with the number of bacteria in the arrangement, and show that the adsorption rates for microcolonies (increasing with exponent ∼1/3) and bacterial chains (increasing with exponent ∼0.5 − 0.8) are substantially lower than for well-mixed bacteria (increasing with exponent 1). We further show that, after infection, the spatially clustered arrangements reduce the effective burst size by more than 50 % and cause substantial superinfections in a very short time interval after phage lysis.

**SIGNIFICANCE:** When bacteria forms clusters they substantially change their exposure to invading phages and other external agents from the well-mixed scenario. Despite this fact, much research has focused on and is focusing on using well-mixed bacteria. Understanding the kinetics of the spatial structures is paramount to developing robust analyses and theories of experimental results. We carefully investigate how the clusters lower the adsorption rate of external phages and how the clustering modifies the hit probabilities for the bacteria.

## INTRODUCTION

The interaction of predators and prey is a widely studied phenomenon and is observed across all scales of life. From wolves and elks to foxes and rabbits all the way down to phages and bacteria. Across all of these taxa, the prey can utilize a plethora of defensive strategies to increase their odds of survival in the presence of their predator. Many of these strategies have evolved to counter a predator in very specific circumstances but some strategies are more general.

In this paper, we explore the benefits of herding from the perspective of bacteria. While bacteria do not form herds in the traditional sense of the word, they often form clusters or aggregates. Herding has been shown to be an effective strategy in some ecosystems, where the localization of prey increases the time it takes for the predator to find the herd of prey (1). There are however also strong negative consequences of herding, e.g. if a herd is found the predator can take down several of the prey at once.

Bacteria and phages constitute a very extreme predator prey system where the phages, upon killing a bacterium, produces on the order of 100 new phages — some times thousands (2). At face value, this extreme amplification of predators should make it impossible for the bacteria to survive, but an ongoing “arms race” between the bacteria and phages have allowed the bacteria to sustain their population. This co-evolution is extensively studied and many intricate bacterial defence mechanism have been identified, such as Restriction-Modification systems(3), CRISPR(4, 5), and abortive infections systems(6).

Bacteria in nature are often found as separated bacteria or as dense arrangements such as microcolonies and biofilms(7–12). This is often due to external circumstances which dictate how the bacteria can be distributed. For example, bacteria cannot move freely in the soil or other solid media. In liquids, the shear forces might prevent the bacteria from forming clusters, but this is not always the case(13). In these environments, the spatial structure has been shown to strongly increase the survival of the bacteria in the presence of phages(14–16), but there is an important aspect of this structure which has not received a lot of study - namely the delaying of phages reaching the bacteria.

When structured, bacteria can associate into various arrangements ranging from simple pairs (diplococcus and diplobacillus) to continuous chains (streptococcus and streptobacillus) to elaborate clusters (e.g. staphylococcus, palisades, and microcolonies). While each arrangement might confer some communal benefits, via quorum sensing, cross-feeding etc., we consider only how such arrangements also will act as a phage defence.

Previously, it has been suggested that the bacterial arrangements have a negative effect on bacterial survival since the larger size of the bacterial arrangement means that the phages are adsorbing at higher rates (17). However, this study fails to take the increased distance between the arrangements into account which leads to net smaller adsorption to bacterial arrangements — compared to if the bacteria were unstructured (similar to the study of(1)).

There are several studies which show that the spatial structure of densely packed bacteria might further reduce the ability of phages to attack the bacteria. For example, it is believed that the extracellular matrix of biofilms strongly captures the phages (14, 18) and that bacterial microcolonies only expose the surface bacteria to the phages and thereby shield the central bacteria from the phages (15).

## MATERIALS AND METHODS

### Simulations

We base our simulations on the code used in Ref. (15). Here, the authors simulated individual spherical bacteria and individual phages. In this work, we expand their model to account for elongated bacteria, which requires some modifications. Now the position of the bacteria is described by two poles 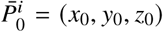 and 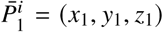. We model the bacteria by the line-segment connecting 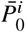 to 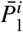. The two poles are considered to be connected by a spring of length *L*, leading to an internal potential of 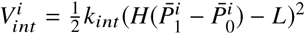, where 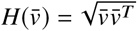 is the length of vector 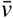. Each bacteria is assumed to have an extent *R* around the central line-segment, and any bacteria that overlaps with this region will be pushed away. We implement this cell-cell repulsion by use of a piece-wise potential. When the cells overlap, we use a spring-like potential, while we have zero potential otherwise. If we define the vector 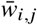 as the shortest vector between the line-segments of two bacteria, the interaction potential takes the form:

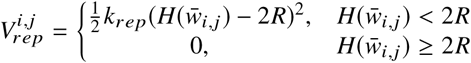

These two potentials are sufficient to describe how elongated bacteria retain their shape and how they interact with their immediate neighbours. For this work, we are interested in modelling chains of bacterial, so we introduce a potential to keep the chain connected:

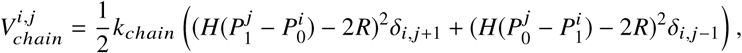

where *δ*_*i,j*_ is the Kronecker delta function. This potential connects the poles of two consecutive bacteria by a spring potential.

In total, the potential felt by cell *i* is

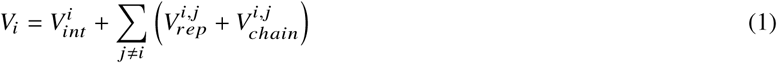

Since we now consider elongated bacteria, we also need to change how hit-detection works for the phages. In the original code, the phage is considered to have hit a bacterium, if its position is less than one cell radii away. In our case, a phage collides with a bacterium if the distance between the phage and the bacterial line-segment is less than *R*.

We generate three types of bacteria arrangements: well-mixed, chains, and microcolonies. For the well-mixed bacteria, the position and orientation of each bacterium are uncorrelated and they are initialized uniformly in the simulation space. In addition, we set the parameter *k*_*chain*_ = 0 N /µm.

When generating the bacterial chains, we do so in an iterative fashion. Starting with one randomly oriented bacterium, we, for each consecutive bacterium, draw a random angle θ_*i*_ between [0, Θ] from a uniform distribution and place the next bacterium a distance 2*R* from the previous and rotated by the angle θ_*i*_ (see figure 1B for illustration). Our chains are thus closely related to the freely-rotating chain model from polymer physics with the exception that our chains will locally repel any overlapping parts. The spherical colonies are generated by first generating a chain with Θ/2π = 0.5 and *L* = 0.1 µm, corresponding to consecutive bacteria on average being at right angles with each other. We then set *k*_*chain*_ = 0 N/µm and expose the bacteria to a force 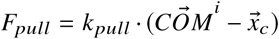 which pulls each bacterial center of mass, 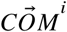, towards the arrangement center 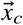. After compressing the arrangement a time *T*_*compress*_ (*N*) (see Table S2 in the supplement for values), we set the cell lengths to the original value *L* and let the bacteria relax. This method, while somewhat cumbersome, generates bacterial arrangements which are very spherical. If, for example, the cell length is not set *L* = 0.1 µm in the beginning, almost all bacteria in the arrangement align to form a disk of bacteria, rather than a sphere.

**Figure 1:**
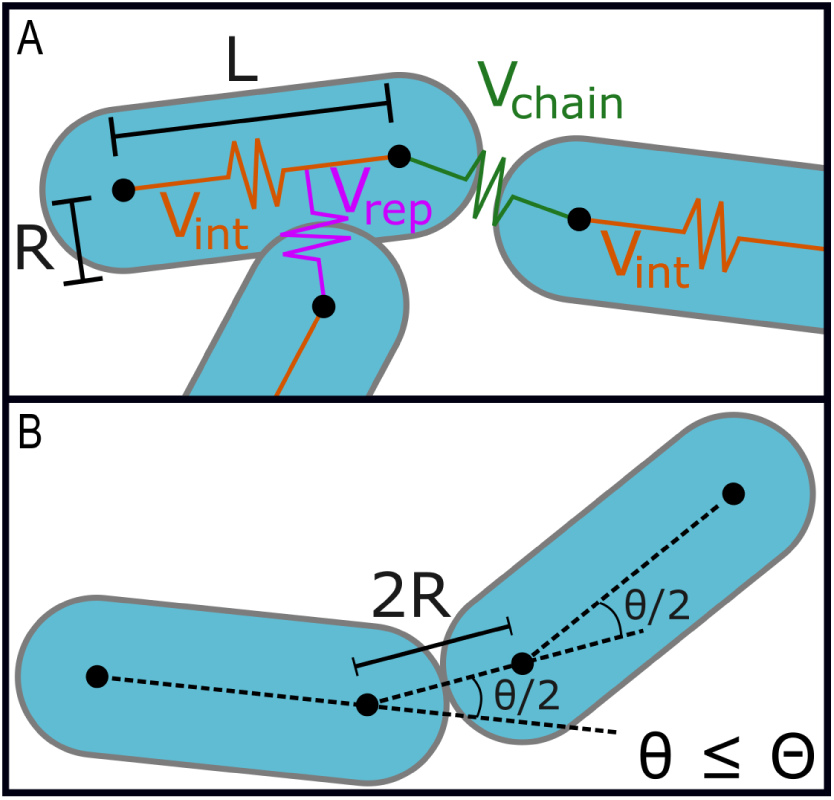
Interaction overview. (A) Illustration of dimensions of the bacteria and the potentials in the simulation. (B) Illustration of a chain being generated.

For each of the bacterial arrangements, there is a reasonable probability that some bacterial pairs overlap, and we, therefore, let the arrangement relax for some time before adding the phage to the simulation. We have operationally chosen to relax the bacteria in intervals of *T* = 0.1 h, after which we measure the maximum overlap, *d*, between any bacterial pairs. If this overlap is more than 1% of *R* we repeat relaxation steps until the overlap drops below this value or until the overlap stops decreasing.

In the supplement, we show that our bacterial arrangements have very little overlap after relaxation (see figure S1 for details).

When adding phages to the system, we must first consider how large of a volume around the bacterial chain we should simulate. For each bacterial chain and spherical colony that we generate, we determine the smallest cube that can encapsulate the bacteria (minimum bounding box) of size *l*_*bb*_ × *l*_*bb*_ × *l*_*bb*_. We then construct a test volume of that is 9 times larger and release test phages inside the volume. We let the phages adsorb for *T* = 0.1 h or until a fraction of 1 − *e*^−1^ phages have been captured to estimate the adsorption rate η. This estimate of η allows us to construct a simulation volume of size *l* × *l* × *l* which is large enough, that 90 % of the phages remain free at time *T* = 1 h. For some simulated chains, the bacterial arrangement is very elongated and the estimated length *l* approaches the length of the bounding box *l*_*bb*_, and we, therefore, impose that *l* must be larger than 2*l*_*bb*_.

Ideally, we would like the phages to be spatially distributed according to the steady-state distribution. To test how robust our phage distribution is, we have developed an algorithm to equilibrate the phage distribution. The algorithm works by simulating phage diffusion for *T*_*eq*_ time where phages, upon hitting a bacterium, is relocated to a random location in the simulation space. In the supplement, we show that our adsorption measurements do not change significantly as we let the phage distribution equilibrate beforehand (see figure S9 for details).

Finally, we need to determine what the time-step used in the simulations should be. We use a chain of size *N* = 10, with Θ/2 = 0.3 π. Using the algorithm above yields a required box size of *l*≈ 180 µm, but we here use a conservative size of *l* = 350 µm. In the supplement, we test several time-steps and find that convergence is achieved around Δ*T* = 10^−6.5^ h and we, therefore, run our simulations with Δ*T* = 10^−7^ h.

We add phages in two ways. For the simulations that measure the adsorption rate and the shielding effects, we uniformly add at least 10^5^ phages to the simulation volume. In the cases where we must impose a volume of length 2*l*_*bb*_, we scale the number of phages so that the density is conserved. That is, we add (2 *l* _*bb*_/*l*)^3^ · 10^5^ phages. When we measure the secondary infections, we remove the selected bacterium from the simulation and introduce 10^4^ phages at its location. These phages are distributed uniformly inside the replaced bacterium and are then free to diffuse in the system.

### Data availability

The code and generated data files are available at the online repository located here(19): https://github.com/RasmusSkytte/BacterialChains/tree/v1.0

## Experiment

We use the Escherichia coli strain, SP427(20) which is derived from MC4100 and encoded with a *P*_*A*1/*O*4/*O*3_::*gfp*mut3b gene cassette(21). This strain was incubated overnight in YT broth (0.8 % W/V Bacto tryptone, 0.5 % W/V NaCl, 0.5 % W/V yeast extract) and subsequently diluted to ∼ 10^3^ CFU/ml in a buffer solution (50 mM CaCl_2_, 25 mM MgCl_2_). 10 µL of the diluted bacteria was then mixed with 0.4 mL of a soft agar consisting of 1 % W/V Bacto tryptone, 0.8 % W/V NaCl, 0.5 % W/V yeast extract, 0.5 % W/V Bacto agar, 0.2 % W/V glucose, 50 mM CaCl_2_, 25 mM MgCl_2_ and 10 mM TRIS. Note, however, that the mixture in the experiment behaved different from normal conditions and was substantially less viscous than duplicate conditions. The reason for this deviation is unknown, but we hypothesize that the soft agar was not fully melted before the bacteria were added.

The mixture was plated onto in one well of a 6-well plate (P06-1.5H-N) from Cellvis (P.O.Box 390959, Mountain View, CA 94039) and incubated for 4 h at 37°C before images were taken. The images capture the green fluorescence signal. For visual clarity, the image has been colour inverted and filtered for extra contrast.

## RESULTS

In figure 2, we show a simple example where we compare phages adsorbing to two separated bacteria (cocci) with adsorption to two joined bacteria (diplococcus).

**Figure 2:**
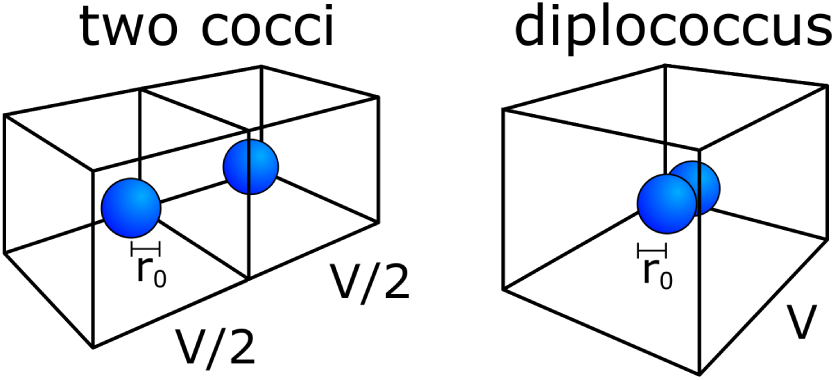
Illustration of two bacterial arrangements. The illustration shows two distinct ways that two bacteria may arrange within a volume *V*. They can be separate, and thus spatially uncorrelated or they can cluster together and form a single arrangement.

Free phages move randomly by diffusion until they encounter a bacterium or they decay. Since their movement is unguided, the time it takes for the phages to reach a bacterial target is well described by diffusion mechanics. Here, we can use a mathematical result derived by Smoluchowski (22), where he shows that the rate of a small diffusing particle hitting a large relatively stationary target is:

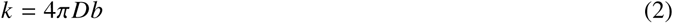

Where *k* is the rate of adsorption, *D* is the diffusion constant for the particles, and *b* is the radius of the target. Notice that *k* has units of m^3^s^−1^. Using this rate, we can compare the difference in adsorption rates for two separated bacteria and the adsorption rate for two joined bacteria. When the bacteria are separated, they each occupy a volume of *V*/2, and we assume they have the same radius of *r*_0_. The phages will then adsorb to either of the bacteria at a rate of:

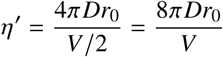

or, since there are two bacteria, the phages will adsorb to *any* bacterium at a rate of:

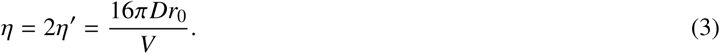

This is the relevant rate since the large burst size makes the finding of subsequent target much faster. If the bacteria are joined together, we will assume they can be treated as a single target of radius *r*_*c*_. This target is assumed to be spherical and has twice the volume of a single bacterium (*V*_*c*_ = 2*V*_0_), and therefore *r*_*c*_ = 2^1/3^*r*_0_. This large target will occupy the full volume *V*, which means that the adsorption rate to the joined target is:

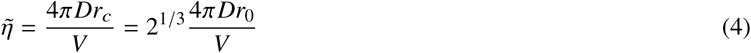

The relative difference in adsorption rate is therefore:

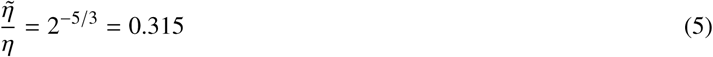

There is then a substantial reduction in the rate of adsorption as when the bacteria are joined together compared to when they are apart. Note that once a phage adsorbs to one of the bacteria in the cluster, the neighbouring bacteria is then almost certainly going to be infected when the other bacterium lyses. In order for this arrangement to have a net benefit, the gain achieved by delaying the initial phage adsorption has to be greater than increased risk to the neighbouring bacteria. Furthermore, the assumption that the joined bacterium can be treated as a spherical target is crude, but this approximation improves as the number of bacteria increases.

As we increase the number of bacteria, the bacteria can form microcolonies, e.g. as in soft agar where they grow to form roughly spherical arrangements (11, 12, 15, 16). Alternatively, the bacteria may arrange themselves in more complicated structures such as biofilms(14, 18, 23) or the bacteria can stick together to form grape-like clusters (staphylococcus), or elongated chain-like clusters (streptococcus/streptobacillus). In an outlier, experiment, we have observed *Escherichia coli* forming a chain-like structure when the soft agar medium was substantially less viscous than expected (see figure 3).

**Figure 3:**
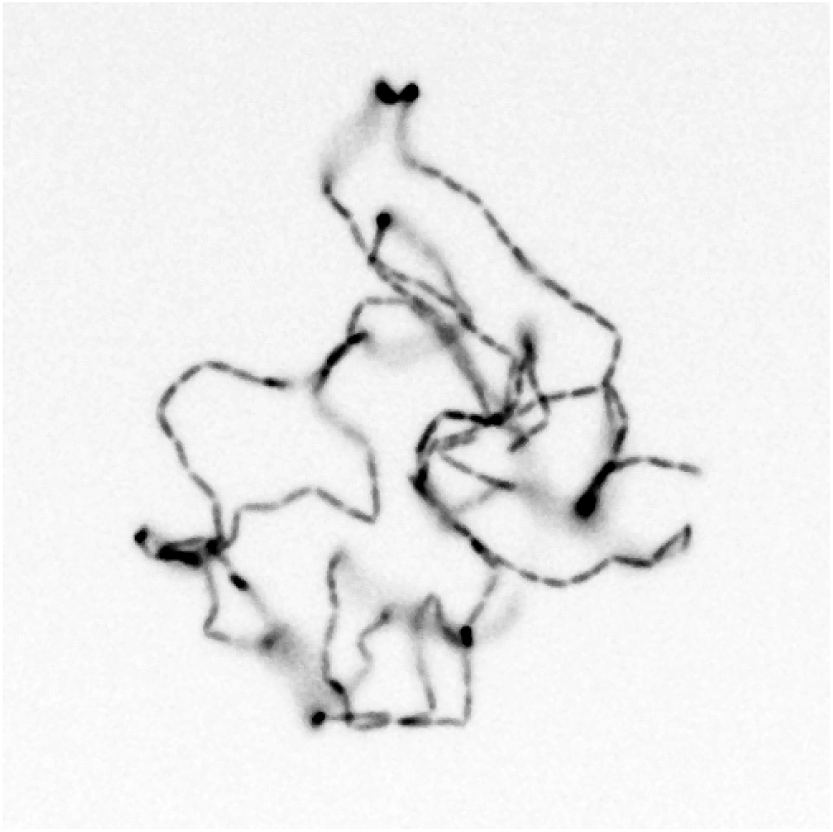
Chain of *Escherichia coli*. In some cases, *Escherichia coli* will grow to form distinct chain-like structures when grown in very soft media. See Materials and Methods for details.

Such chains of bacteria are interesting because the geometry is non-trivial and, depending on the angle between consecutive cells, will have widely varying volumetric scaling. In addition, the dense clusters of bacteria, e.g. microcolonies and biofilm are well studied, but we have found few published studies of the chain-like clusters.

For chains of bacteria, we cannot in general approximate the radius of the bacterial arrangement to be *r*_*c*_ = *n*^1/3^*r*_0_ as we did above for the dense arrangement. Such a chain can either be bunched up, in which case we can approximate with a spherical arrangement, or the chain can be more dispersed, with the most extreme configuration being a straight line.

Using detailed simulations of single bacteria, we construct three types of bacterial arrangements. We test the unstructured arrangement where the bacteria are widely separated (i.e. well-mixed). For structured arrangements, we generate densely packed spherical colonies and elongated chains of bacteria. We generate each arrangement of various sizes *N* and for the chains we use different maximum internal angles Θ, creating chains ranging from (almost) straight lines to small clusters. In figure 4, we show three examples of simulated chains, each with different average internal angles, and one example of a spherical colony.

**Figure 4:**
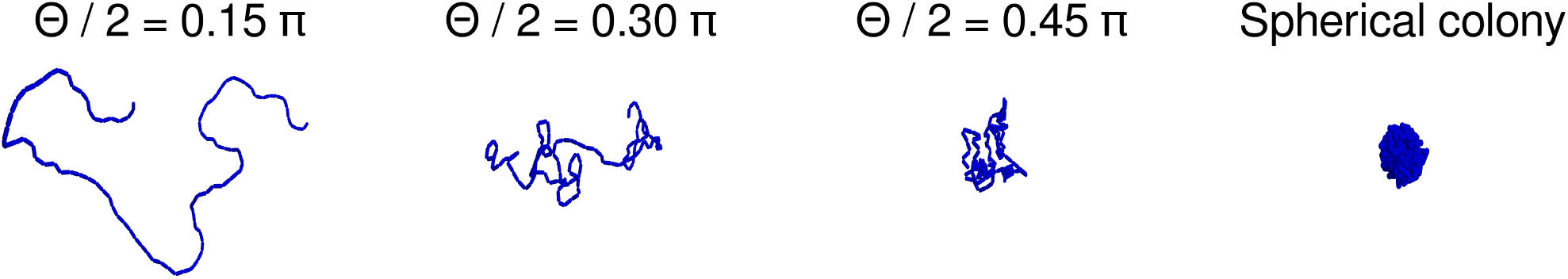
Examples of simulated arrangements. We show examples of chains for three different average internal angles parameters Θ, and one example of a spherical colony. All examples have consist of 100 bacteria.

For each arrangement we generate, we estimate the required volume to allow for 10% phage adsorption within a 1 h window. Short curled-up chains could be simulated by considering relatively small volumes, while long, dispersed chains require a simulation that includes a larger volume for the diffusing phages. We then fill the simulation space with individual phages particles which undergo diffusion until they encounter the bacteria, at which point it is removed from the simulation.

In figure 5A, we show the adsorption rate as a function of the number of bacteria for the various bacterial arrangements. The well-mixed bacteria and the spherical colony effectively constitute the largest and smallest phage targets respectively, with the bacterial chains having intermediate adsorption rates dependent on the internal angle parameter Θ. We fit each family of curves to the function η(*N*) = *k N*^γ^ and find that the exponent γ decreases with the the internal angle parameter Θ (see figure 5B).

**Figure 5:**
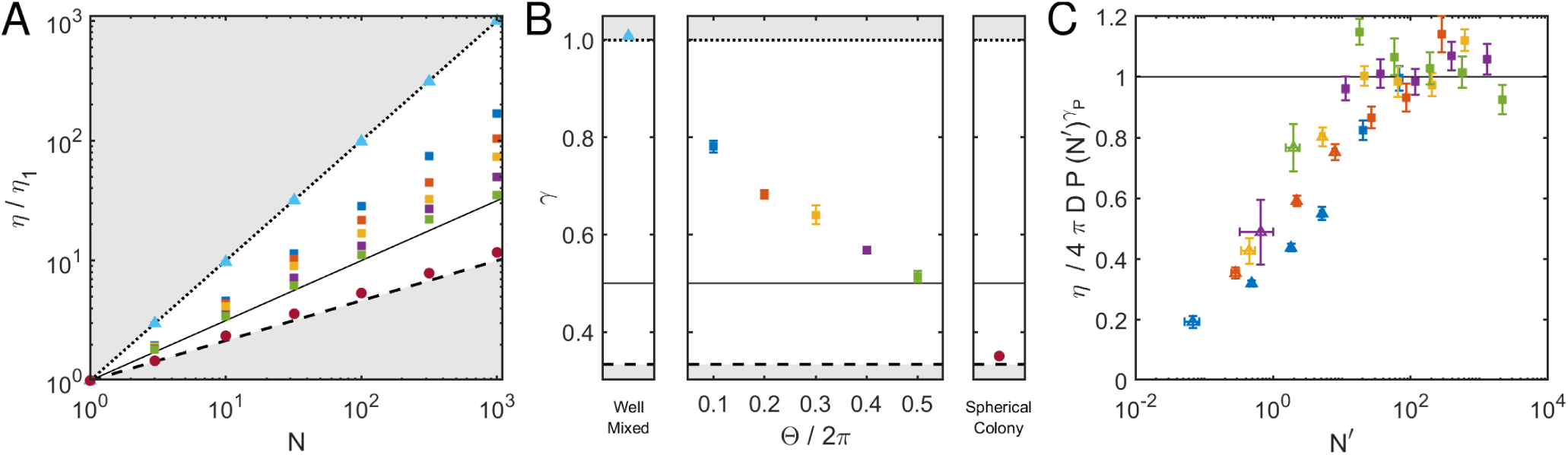
Phage adsorption rate. (A) We show how the adsorption rate scales with the number of bacteria, *N*, in various arrangements. The adsorption rate is highest for well-mixed bacteria (triangles), intermediate for chains of bacteria (squares), and minimal for spherical colonies (circles). The chains have varying values for Θ/2π: 0.1 (blue), 0.2 (red), 0.3 (yellow), 0.4 (purple) and 0.5 (green). Points are error bars showing standard error (not visible). (B) We fit each family of adsorption rates to η = *k N*^γ^ and we show the value of the exponent γ. Error bars indicate standard deviation. (A-B) For comparison, we show three models of adsorption rate scaling: well-mixed scaling η/η_1_ = *N* (dotted line), Random walk scaling η/η_1_ = *N*^0.5^ (dot-dashed line), and spherical scaling η/η_1_ = *N*^1/3^ (dashed line). Grey areas indicate inaccessible areas. (C) A fit to the adsorption rate as a function of *N*, the persistence length *P*, and the phage diffusion constant *D*. Triangles indicate points where *N*_*i*_ < *P*_*i*_. Fitted value is γ_*P*_ = (0.57 ± 0.01) with a reduced χ^2^ of 4.6.

Another way of describing the family of bacterial chains is by the persistence length, i.e. a measure of the stiffness of the chain. We compute the average persistence length for the data-set, *P* (Θ, *N*) across 100 replicates, and compute the effective length *N*^′^ = *N*/*P*(Θ, *N*) of the chains.

The adsorption rate to each family of chains can be modelled by the following equation:

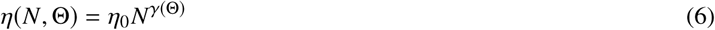

With the measured persistence lengths, we can re-scale all the chains by their persistence length to chains that scale with the same exponent γ_*P*_:

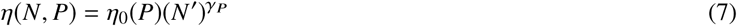

This equation considers the chains to consist of fewer but longer links. With our parameters, this base adsorption rate can be approximated by η_0_ (*P*) = 4π*DP*.

In figure 5C, we then plot the adsorption rate as a function of *N*, and see that for sufficiently long chains these collapse onto a scaling of 4π*DP* (*N* ^′^) γ*P*. We can only reliably estimate the persistence length when the length of the chain *N* is larger than the persistence length, and as a result, we observe deviation from this scaling at small *N* ^′^.

In addition to the reduced adsorption rates of the structured bacterial arrangements, the arrangements non-uniformly expose the bacteria to the surroundings. To quantify this, we track the individual phage collisions and measure the ratio of hits between the most exposed bacterium and the least exposed bacterium. In figure 6, we show how this ratio changes for the bacterial arrangements as the number of bacteria increase.

**Figure 6:**
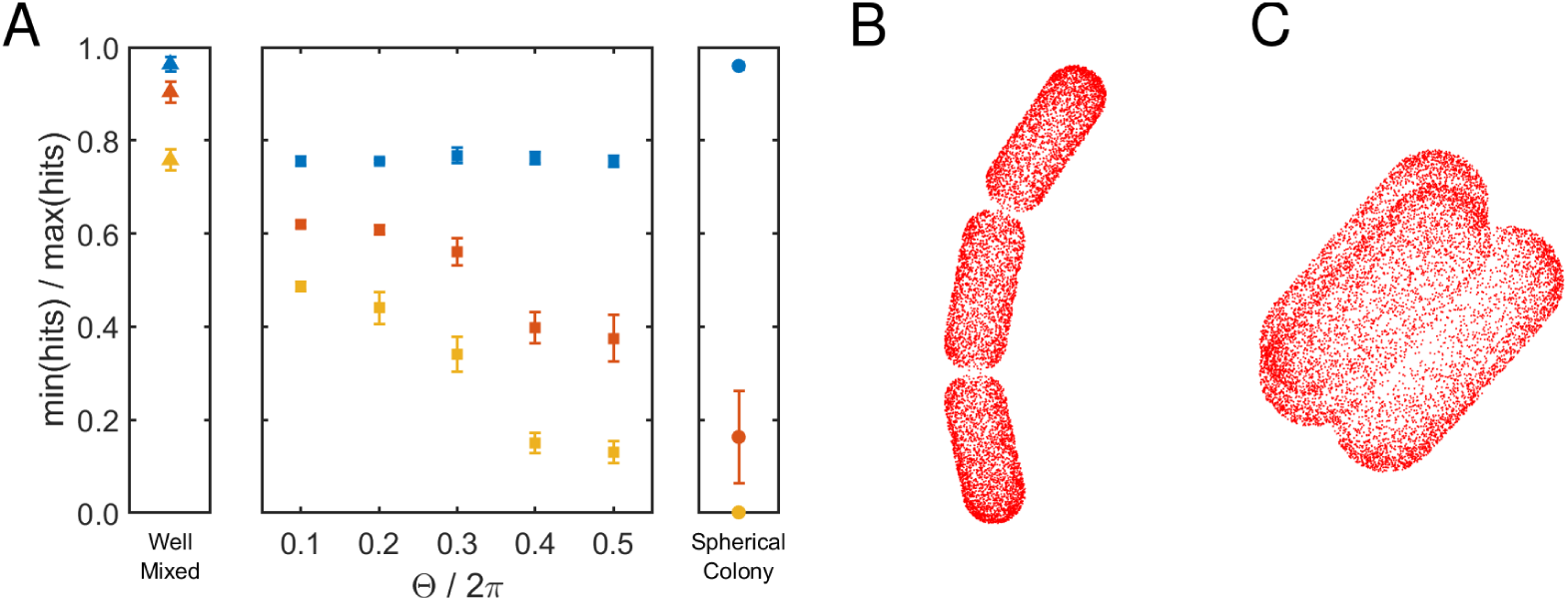
Shielding effects. (A) The ratio of hits between the most exposed bacterium and the least exposed bacterium. The error bars show the measured ratios for arrangements of size *N* = 3 (blue), *N* = 10 (red), and *N* = 32 (yellow). The chain arrangements show increasing shielding with increasing size, *N*, as for the well-mixed bacteria, but also with increasing internal angles Θ when the chains are *N* ≥ 10. The spherical colony can almost fully shield some bacteria when *N* ≥ 10 but exhibit almost no shielding at *N* = 3. (B) Phage hits on a chain of length *N* = 3 and Θ/2π = 0.1. (C) Phage hits on a spherical colony of size *N* = 3.

The well-mixed bacteria show increasing shielding with increasing *N* — However, notice that we take the ratio of the extremal values which will increase as we increase the number of bacteria, even when there is no statistical difference. Even so, the structured arrangements exhibit substantially more heterogeneous phage adsorption with some bacteria experiencing relatively few hits. The spherical colony is especially efficient in shielding bacteria. Already for *N* = 10 some bacteria are almost fully protected against the phage. For *N* = 3, the symmetry of the spherical arrangement means that all bacteria have an equal exposed area, and therefore the phage adsorption is homogeneous (see figure 6C). This is not so for the chain arrangements. Here, the phage adsorption is substantially more heterogeneous even at *N* = 3. One reason for this is that the centre bacteria has two areas of contact with other bacteria, while the end bacteria have only a single area of contact each. Therefore, the centre bacterium exposes a smaller surface to outside and consequently is less likely to be hit by the phages (see figure 6B). As the number of bacteria increase, the chains with larger internal angles begin to outperform the bacteria with smaller internal angles. The smaller internal angles mean that the bacterial arrangement only gets shielding from the difference in contact areas, while the chains with increasing internal angles receive a corresponding increase in shielding as the chains contract and begin to form a spherical colony.

The above results show that 1) the bacterial arrangements delay the phage from reaching the bacteria and 2) the bacteria are not uniformly exposed to the phage attack.

Once found by a phage, the colony is most certainly doomed as the progeny phage will be released immediately adjacent to the remaining bacteria. We next investigate these secondary infections.

For each bacterial arrangement, we do 5 simulations where we lyse a bacterium (3 simulations when *N* = 3). These bacteria are chosen equidistantly along the arrangement. The chosen bacteria are removed from the simulation and 10^4^ phages are spawned in their location. From these phage fates, we bootstrap *β* phages 1000 times, and quantify the secondary infection by measuring 1) the fraction of the *β* phages that remain free 1 hour after lysis, and 2) the fraction of the *β* phages that result in new infected bacteria. These together constitute the effective burst size which we plot in figure 7A. Each of the lysed bacteria has differing probabilities of being hit by an external phage, so we weigh the effective burst size by these probabilities. E.g. a bacterium at the core of a spherical colony has almost zero chance of being hit and should, therefore, contribute little to the result.

**Figure 7:**
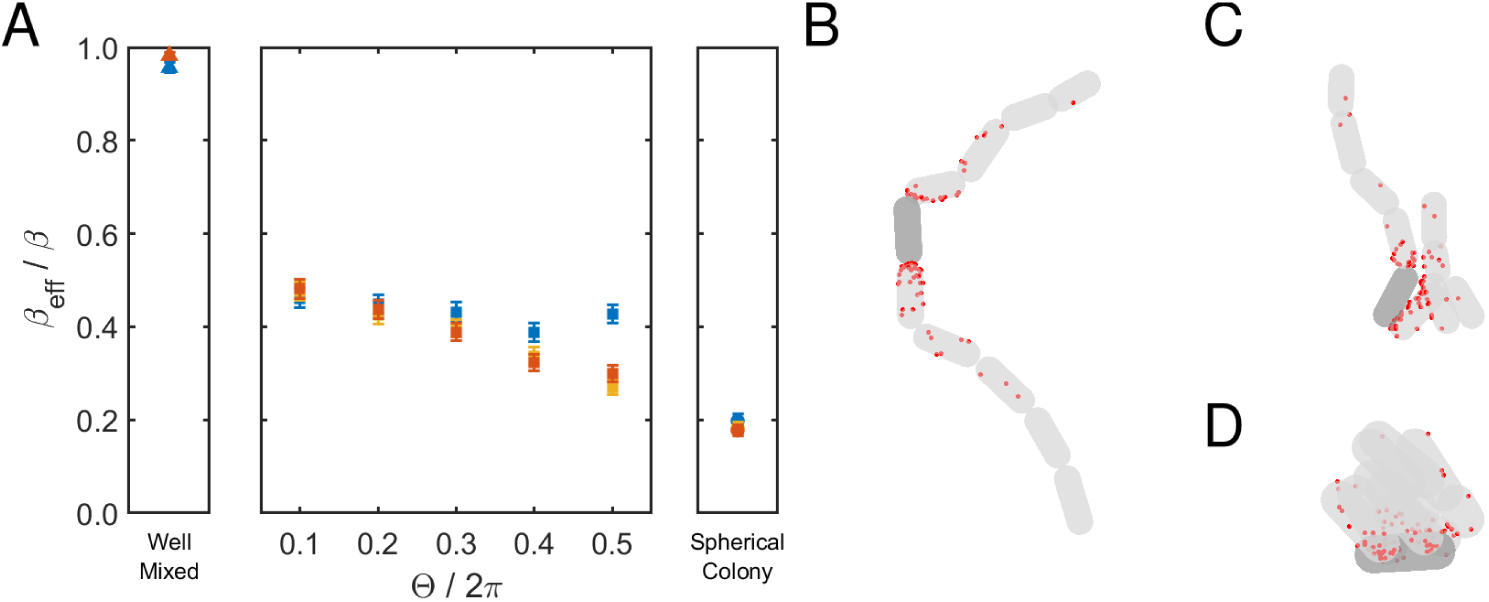
Secondary infections. We measure the effects of phage mediated lysis within the bacterial arrangements. (A) The effective burst size *β*_*eff*_ for *β* = 100, is the number of phages which either cause new infections or remain free to cause future infections (measured 1 h after release). Blue, yellow, red points indicate *N* = 10, *N* = 32, and *N* = 100 respectively. (B-D) Examples of simulated lysis (*N* = 10, *β* = 100). Red dots indicate locations of phage encounters. Dark grey bacterium indicate the lysed bacterium. (B) Chain with Θ/2π = 0.1. (C) Chain with Θ/2π = 0.5. (D) Spherical arrangement.

The effective burst size for the structured arrangements is only a fraction of the case when bacteria are well-mixed. In figure 7(B-D) we show examples of phage collisions from a bootstrapped sample. This highlight how the secondary phages infect the bacterial arrangements non-uniformly.

This non-uniformity means that many phages will be superinfecting the nearby bacteria and thus “wasted”. If the phage is temperate, the increased multiplicity-of-infection (MOI) at the early infection state will cause a significant fraction of the phage to go lysogenic, and thus further reducing the virulence of the phage.

Using the bootstrapped samples from above, we quantify the fraction of infected bacteria that have been superinfected for *β* = 100 (see figure 8A). Note that this measure is sensitive to the chosen *β* value.

**Figure 8:**
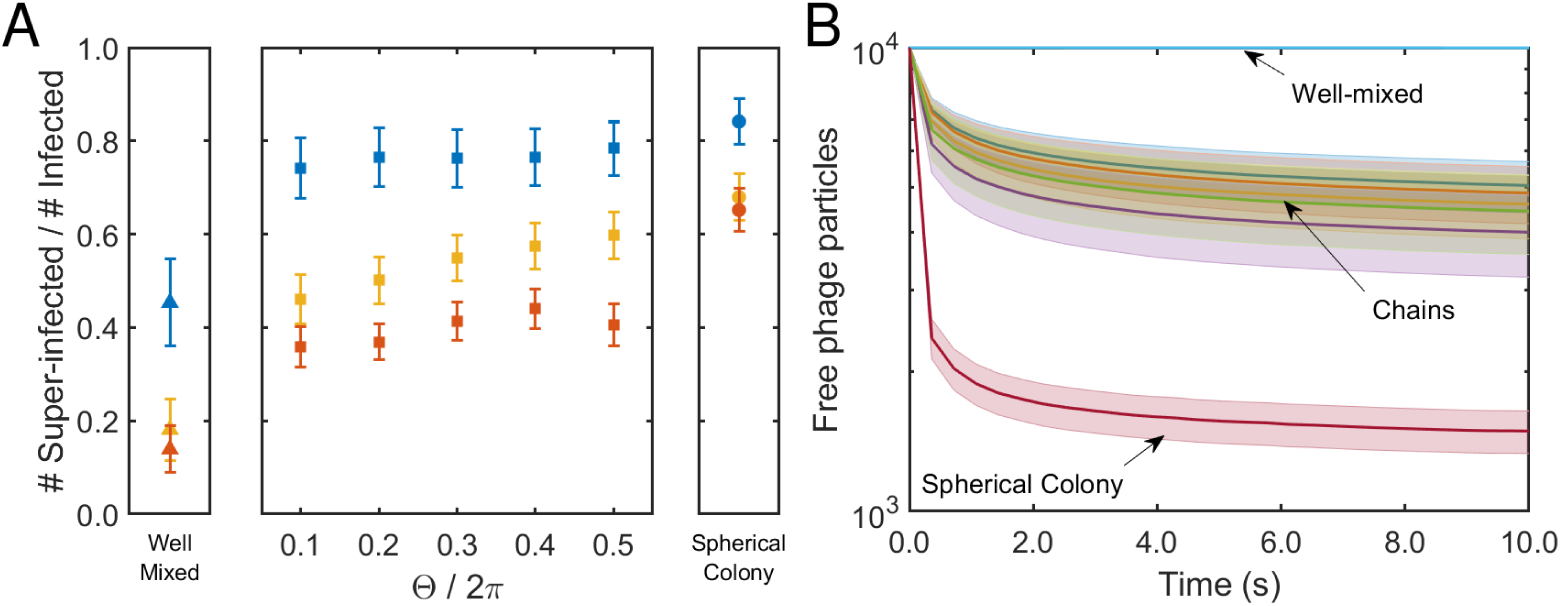
Superinfections. Neighbouring bacteria experience high phage pressure after lysis events. (A) The structured bacterial arrangements exhibit a much higher rate of superinfections than the well-mixed bacteria. Blue, yellow, red points indicate *N* = 10, *N* = 32, and *N* = 100 respectively. (B) Average adsorption curves for *N* = 10. The structured bacterial arrangements show substantial phage capture within a small time frame. Shaded areas indicate standard error of the weighted mean.

In the structured arrangements, the phages are substantially more likely to hit an already infected bacterium than they are to hit an uninfected bacterium. Even when the arrangement contains 100 members, roughly half of the infected bacteria are hit by more than one phage. If the phage is temperate, this suggests that many of the infections will lead to lysogenic bacteria(24).

Furthermore, in these structured arrangements, the progeny phages are released immediately next to susceptible hosts and when considering the time-scale for adsorption (see figure 8B) the phages are adsorbed within a very small time frame, further increasing the probability for the bacterium become lysogenic.

## DISCUSSION AND CONCLUSION

Our investigation shows that the clustering of bacteria substantially changes to their exposure to invading phages. This is a result of several factors: 1) The clustering of bacteria increases the time it takes for the phages to locate the bacteria. 2) The heterogeneous distribution of bacteria within the clusters increase the likelihood of phages superinfecting bacteria. 3) The phages released by lysis are very likely to superinfect bacteria causing the effective burst-size to decrease substantially.

These changes have profound effects on the virulence of the phage and the effectiveness of bacterial defense systems. For example, if the phage is temperate, the short window of time after lysis where the bulk of the superinfections occur are likely to cause a high fraction of lysogenic infections which reduces the virulence of the phage attack. The high fraction of superinfections also modifies the effectiveness of some bacterial defence mechanism, such as abortive infections systems (6), which in this spatial context can be used to negate the effect of several phages at once. The high local MOI may also protect the bacterial populations by causing “lysis from without” (25, 26), whereby the phages themselves cause an aborted infection. Interestingly, these effects are present even at very small clusters, e.g. consisting of three bacteria. It is worth keeping in mind that the increased MOI stemming from the structure of the bacterial cluster comes in addition to increase in MOI that spatial structure alone brings(16, 27).

Similarly, some pages, like phage T4, exhibit lysis inhibition whereby they delay the onset of lysis if the MOI high (28, 29), and such phages are thus much more likely to enter this state when the bacteria are structured than when they are well-mixed.

When bacterial arrangements become large, the bacteria begin to compete internally for resources (12), and this competition may out-weight the benefits of the phage defence described above. This suggests that short to medium-length chains of bacteria and small bacterial colonies are benefiting from the phage defence properties of their spatial structure without a substantial reduction in available resources.

The heterogeneity in resource availability and phage exposure suggests that the bacteria on the edge of the bacterial clusters, due to the higher phage presence, should invest more in bacterial defenses while the bacteria closer to centre, due to the reduce nutrient level, should invest more in growth.

Several other papers have shown that highly structured bacterial arrangements are capable of protecting the bacteria such as when they form a biofilm (14, 18) or when they form microcolonies (15, 16). Our results show that even small, weakly structured bacterial arrangements can cause substantial changes to the interaction with phages which may help explain why bacterial clustering is so prevalent even in liquid conditions (13).

There are several documented synergistic mechanism for bacteria living in clusters: Bacteria may utilize quorum sensing, whereby bacteria modify their local environment when the local population density is high, to share the burden of resource utilization (30), or the bacteria can evolve additional phage defences such as abortive infection systems (6). The reduced phage pressure and the benefits of group living from existing as clusters, provide the bacteria with a “win-win” scenario and may explain why bacteria have evolved mechanisms to remain clustered even in liquid environments.

## AUTHOR CONTRIBUTIONS

All authors designed the research. R. S. E. carried out all simulations and analyzed the data. All authors wrote the article.

## ACKNOWLEDGMENTS

The authors thank Jannik Vindeløv for helpful discussions and Frej Andreas Nøhr Larsen for his help in conducting the experiment.

This project has received funding from the European Research Council (ERC) under the European Union’s Horizon 2020 research and innovation program under grant agreement No. 740704.

## SUPPORTING CITATIONS

References (31, 32) appear in the Supporting Material.

